# A Comparative Study of MBTI and Learning Style-Based Grouping for Enhancing Group Effectiveness and Balance in a Pedagogical Setting

**DOI:** 10.64898/2026.07.05.736636

**Authors:** Basith Nasik, Shanna Nifoussi

## Abstract

Effective group work is central to Problem-Based Learning (PBL) in higher education, yet the optimal strategy for forming student groups remains unclear. This study compared MBTI-based grouping, informed by personality types and Keirsey temperaments, with Learning Style–Based (LSB) grouping, grounded in Kolb’s Experiential Learning Theory, to assess their impact on group functioning and role performance. Participants were undergraduate students enrolled in Cell Biology (Fall 2022 and Fall 2023) and Introduction to Biology Laboratory (Fall 2023) courses. Students completed MBTI and Kolb Learning Style assessments, and groups and roles (Leader, Communicator, Organizer) were assigned accordingly. Results indicated that LSB-based groups consistently outperformed MBTI-based groups across multiple performance metrics, including productivity, listening, sense of safety, belonging, and overall satisfaction. All metrics showed statistically significant decreases in MBTI-based groups except contribution, which did not differ significantly between grouping strategies. Role performance ratings were significantly higher for Leaders and Communicators in LSB groups, while no significant differences were observed for the Organizer role. Correlation analyses revealed that satisfaction was strongly associated with perceived productivity in MBTI-based groups, whereas in LSB-based groups, satisfaction was more strongly correlated with psychological safety. These findings suggest that learning style alignment may better support effective collaboration and group climate in PBL settings than personality-based grouping.

## INTRODUCTION

Over the past five decades, the aims of higher education have shifted from a traditional emphasis on knowledge for its own sake to a more pragmatic focus on career readiness and the development of transferable skills.^1^As universities expand curricula to span disciplines and align with workforce needs, they increasingly prioritize competencies such as communication, critical thinking, collaboration and adaptability.^2^ This evolution reflects broader societal changes such as economic pressures, technological advances, and rising expectations that higher education should contribute directly to employability. Consequently, universities are reimagining their role as launchpads for professional success, alongside their core mission of learning.^3^

In today’s educational settings, group-based learning has gained prominence, particularly through approaches such as Problem-Based Learning (PBL). A persistent challenge, however, is forming balanced and effective groups that leverage individual members’ strengths. A balanced and effective group is one in which the composition and interaction of the group members allow for the achievement of the group goals, frequently oriented towards problem solving. Personality types (e.g., Myers-Briggs Type Indicator-MBTI) and learning styles (e.g., Kolb’s learning styles) are two widely studied frameworks that offer insight into how individuals interact, process information, and contribute to group work.^4,5^

The Myers-Briggs Type Indicator (MBTI), developed in the early 1940s by Isabel Briggs Myers and Katharine Cook Briggs, is a self-report questionnaire that is widely used to categorize respondents into 16 personality types, defined by preferences on four dichotomous dimensions: Introversion/Extroversion, Sensing/Intuition, Thinking/Feeling, and Judging/Perceiving.^6^ Using MBTI in educational settings can offer tremendous opportunities for creating effective and balanced groups. Instructors can utilize the personality types of their students to better understand distinct communication styles and their ability to collaborate, thus fostering an inclusive and supportive classroom environment. By understanding individuals’ personality types makes team building more productive and provides the opportunity to create small learning environments where individuals can apply their strengths in a synergistic manner to achieve the common goal of group tasks. Overall, resulting in greater productivity and satisfaction among the individual members in the group.

Several years after MBTI was created, in 1984 David A. Kolb introduced his Experiential Learning Theory (ELT), inspired by the work of the gestalt psychologist Kurt Lewin, as well as John Dewey and Jean Piaget.^7^ELT articulates a four-stage learning cycle (Concrete Experience, Reflective Observation, Abstract Conceptualization, Active Experimentation) and four corresponding learning styles. ELT emphasizes that individuals favor different modalities of learning and that teaching approaches should accommodate these variations to enhance engagement and learning outcomes.^7^The pedagogical implication is that providing opportunities to engage multiple modalities can make learning more interactive and effective for diverse learners. Learning styles, by definition, are the different ways in which individuals learn, process, and recall information. For example, some learners may prefer visual stimuli in the form of diagrams and charts, whereas others may learn optimally through auditory stimuli or practical experience (e.g., kinesthetic).^8^ This led to the pedagogical approach that individualizing teaching practices is crucial to meet the various learning style demands of the learners. By doing this, instructors can make learning more interactive and productive, thus leading to increased student productivity and satisfaction.

### MBTI and Learning in Educational Settings

MBTI offers a structured lens through which to understand individual differences in behavior, emotional processing, and communication.^9^In the classroom, MBTI can inform how students prefer to learn and interact.^10^For example, Extraverts may thrive in discussion-heavy group settings, while Introverts may prefer reflective or independent tasks. Thinkers value logic and structure, whereas Feelers may prioritize emotional connection and harmony in group discussions. By identifying these preferences, instructors can design learning groups that engage each students’ strengths, encourage complementary collaboration, and reduce potential friction. Furthermore, MBTI can help identify group roles (e.g., Facilitator, Recorder, Leader) based on a student’s natural inclinations.

### Kolb’s Experiential Learning Theory and Learning Styles

Kolb’s ELT emphasizes that individuals demonstrate consistent preferences in how they perceive and process information. Rather than focusing on the underlying learning cycle, the model is often applied through its four learning styles, which describe how learners approach tasks, solve problems, and collaborate in group settings.^7^

- Divergers: Excel at viewing situations from multiple perspectives.
- Assimilators: Prefer structured understanding through conceptual models.
- Convergers: Practical problem-solvers who apply ideas to real-world scenarios.
- Accommodators: Learn by doing, favoring hands-on experience over theory.

These learning styles translate into distinct preferences for information intake and output—visual, auditory, kinesthetic, and reflective modalities. Matching instructional methods and group tasks to these preferences has been shown to improve engagement and academic outcomes.^11^

While MBTI and learning styles are different frameworks, they are not mutually exclusive. Personality can influence learning style; for instance, extraverts (E) usually prefer active or kinesthetic learning experiences, while introverts (I) usually align more with reflective or solitary learning experiences.^12^ Similarly, judging (J) MBTI types lean towards structured, goal-driven group activities, while Perceiving (P) MBTI types excel in a flexible, exploratory learning environment.^13^

### Problem-Based Learning (PBL) and the Role of Grouping

Problem-Based Learning (PBL) is an educational pedagogy used regularly in medical schools, that immerses students in solving real-world, open-ended problems. It fosters critical thinking, creativity, and cooperative learning.^14^Group work is central to PBL, requiring students to share knowledge, challenge assumptions, and co-develop solutions. Therefore, the composition of these groups is crucial. Effective group composition can: promote diverse perspectives, improve decision-making, enhance problem solving efficiency, prevent groupthink, and overall maximize individual contributions. However, the challenge lies in understanding which individual factors, personality or learning style, better predict successful group dynamics and outcomes.^15^

Previous work has shown that forming small learning groups based on MBTI facilitates smoother interpersonal dynamics.^16^ Utilizing MBTI indicators allows an instructor to predict a students’: Leadership emergence (e.g., ENTJs), Communication styles (e.g., F types may mediate conflict), and Problem-solving preferences (e.g., T types for logic-based tasks).^16^However, MBTI also have some notable limitations when compared to more empirically grounded models such as the Big Five. The Big Five personality model, also referred to as the Five-Factor Model (FFM), conceptualizes personality along five continuous, evidence-based dimensions: Openness to Experience, Conscientiousness, Extraversion, Agreeableness, and Neuroticism.^17^ Based on this limitation, it is a valid claim to make that MBTI may not fully account for how individuals *learn*, only how they *interact*.^18^

Despite the advantages of student learning, group learning is not universally effective. Some personality types, e.g., introverts, or reflective processors, may find group settings overwhelming or inefficient. Similarly, mismatched learning styles can lead to communication breakdowns or unequal participation. For example: a reflective learner/introvert may withdraw during rapid brainstorming, an active learner/extravert may dominate discussions without realizing others’ discomfort, a kinesthetic learner may become disengaged in discussion-heavy settings.^19^Understanding and perhaps anticipating these dynamics is essential for optimizing group-based learning outcomes. This paper aims to address the research question, which of the two frameworks, MBTI vs Learning Style, is a better indicator for the formation of balanced and effective groups in an educational setting.

## METHODS AND PROCEDURES

### Participants

Participants of this study were University of Wisconsin-Superior undergraduate students enrolled in Cell Biology (Fall 2022 and Fall 2023) and Introduction to Biology Lab (Fall 2023, Section 1 and 2). Students were invited to participate in the study on the first day of class. Specifically, students were informed that the course would heavily utilize small group work, and that their participation in the study was voluntary and would not impact their group assignment grades nor their overall course grade.

### Student Information Survey

Students enrolled in each course associated with study were asked to complete a ‘student information survey’ which asked about their goals for the class, their strengths and weaknesses in the subject matter, their past experience working in small groups, and their preferred role in a small group (e.g., leader, communicator, or organizer). Additionally, students were asked to complete two online self-assessments, Kolb’s Learning style inventory test, and a MBTI personality assessment, and provide the results. The information from this survey was used to form small groups and assign students to respective group roles (i.e., Leader, Communicator, Organizer).

### Group Formation and Role Determination

The MBTI-based groups were grouped according to the Keirsey Temperament Sorter, a framework derived from the MBTI **(Figure 1)**. This framework organizes the 16 MBTI into four broader temperaments: Guardian (SJ), Artisan (SP), Idealist (NF), and Rational (NT) based on shared behavioral patterns, motivations, and communication styles.^20^

**Figure 1.**
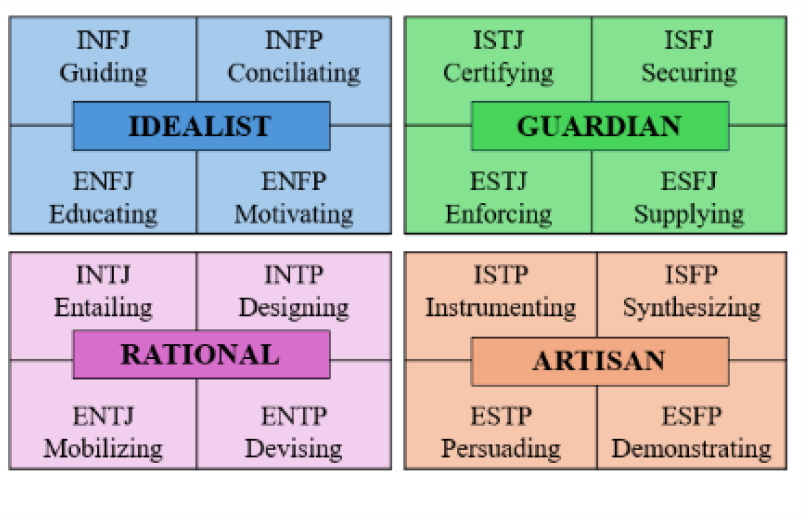
Summary of the Keirsey Temperament Sorter Framework used. Represents the overview of the four Keirsey Temperament categories-Idealist, Guardian, Rational, and Artisan.^20^

- Guardian (SJ): Practical, Detail-oriented, and duty-driven individuals who value structure, stability and responsibility.; they tend to excel at organizing, maintaining order, and ensuring reliability within groups.
- Artisans (SP): Spontaneous, action-oriented, and adaptable, often thriving in hands-on environments that require improvisation, creativity, and quick decision making; they bring flexibility and energy to collaborative setting.
- Idealists (NF): Empathetic, Imaginative, and value-driven, focusing on personal growth, harmony, and meaningful relationships; they are typically strong motivators and communicators who seek to inspire and connect with others.
- Rationals (NT): Analytical, strategic, and independent thinkers who prioritize logic, efficiency, and problem-solving; they tend to approach challenges systemically and excel at conceptual reasoning and innovation.

Based on their MBTI, the students were first classified into their corresponding Keirsey temperament category. Students were then grouped-using a stratified-composition approach, where each group was intentionally formed to include one representative from each of the temperaments. This was done to ensure balanced representation of different temperament types within each group. Typically, each group consisted of three to four students.

Once the groups were made, the individuals in the groups were assigned specific roles (Leaders, Communicators, and Organizers) based on their MBTI **(Table 1)**. Each group had only 1 leader and 1 communicator, and 1-2 organizers.

**Table 1.** MBTI personality types grouped into functional roles for PBL: Leaders (direction and accountability), Communicators (dialogue and connection), and Organizers (detail and quality control).^21^.

| GROUP ROLE | MBTI TYPES | RATIONALE |
| --- | --- | --- |
| **Leaders** | ENTJ, ESTJ, ENFJ, ESTP, ENTP | These types are natural at setting direction and structure, motivating others, and ensuring progress by keeping the group accountable to timelines and goals |
| **Communicators** | ENFP, ESFP, ESFJ, INFP, INFJ | These types excel at connecting with others, expressing group ideas clearly, and serving as bridges between the team and external stakeholders |
| **Organizers** | INTJ, INTP, ISTJ, ISFJ, ISTP, ISFP | These types thrive in analyzing details, refining outputs, and ensuring the final product is both accurate and high-quality, keeping the project grounded and complete. |

Following group formation and role determination, each student was emailed to notify them of the role they were assigned, a detailed explanation as to how their personality assessment contributed to the role assigned, and the expectation of their role in their group **(Table 2)**.

**Table 2.** Group roles and expectations in a PBL setting: Leaders (planning, deadlines, and accountability), Communicators (spokesperson and assignment submission), and Organizers (quality control, finalization, and idea contribution).

| GROUP ROLE | EXPECTATIONS |
| --- | --- |
| **Leaders** | Establish a clear timeline and plan for all the group activities, responsible for establishing deadline for the final project (human bio course only), Ensuring the work is evenly distributed and everyone in the group is participating in and contributing to the group activity, Ensures the group stays on task and the assignment is being completed in a timely manner. |
| **Communicators** | Serves as a group spokesperson checking in with the instructor when clarification is needed, submitting group assignments and participating on behalf of the group on class discussion boards. |
| **Organizers** | Quality control and finalization, overseeing the finalization of the infographic (human bio only), reviewing the content, design and overall quality of all the assignments the group is submitting. Contribute new ideas and perspectives to the group. |

In contrast, groups formed based on learning style assessments (Kolb’s) were organized homogeneously: students identified as *active* learners were grouped together, while *reflective* learners were placed in separate groups. Care was taken to ensure that no group contained a mix of active and reflective learners. Additionally, students’ role preferences (e.g., leader, co-leader, organizer) were considered during group formation to ensure a balanced distribution of roles within each group.

### Group Evaluation

Throughout the course, participants were tasked with engaging in various collaborative activities, such as online discussions and problem-based learning (PBL) assignments. Students in Cell Biology were asked to assess their group learning environment throughout the course at the end of each structured unit with exam wrappers. Exam wrappers included 10 questions answered on a 5-point Likert-scale with two open-ended questions. In addition to this, at the end of the course participants enrolled in ALL courses involved in the study were asked to assess their group learning environment and role performance through completion of course wrappers. Course wrappers also included 10, Likert-scale questions in addition to three Likert-scale questions pertaining to role performance of each assigned group role (e.g., Leader, Communication, Organizer). The exam and course wrappers included questions such as: “I feel that my role in the group was clearly defined,” and “I believe that the group worked well together to accomplish the project.” Students were asked to respond to the wrapper questions to assess the extent to which participants felt that the group dynamics and role fulfillment contributed to the success of the project using a 5-point Likert scale where 1 represented Strongly Disagree, and 5 represented Strongly Agree. The data collected from these wrappers was used to assess group performance, satisfaction, and the effectiveness of role assignments in each group type, and the overall data was used to draw comparisons between MBTI and Learning style-based groups.

### Data Analysis

SPSS analytical software and Microsoft Excel spreadsheets were used to analyze all the data. To find out if there were any statistically significant variations between the two group types in terms of role fulfillment, group effectiveness, and general satisfaction, t-tests were used. To ascertain the connections between individual group dynamics and group performance results, Pearson’s correlation was used to evaluate correlations between group variables (e.g., group satisfaction, productivity, and role fulfillment).

### IRB Permissions’ number

All study procedures involving human participants were reviewed and approved by the University of Wisconsin– Superior Institutional Review Board (IRB approval number E159).

## RESULTS

### Demographic of Participants

In Fall 2022, 15 students were enrolled in the Cell Biology course, and all were assigned to LSB-based groups. In Fall 2023, 21 students were enrolled in Cell Biology, and all were assigned to MBTI-based groups. For the MBTI population, in relevance to the ‘David Keirsey’s Temperament’, 9 students fell under the ‘Guardian’ category, 6 ‘Idealist’, 6 ‘Rational’ and 0 ‘Artisan’. Based on their MBTI, 8 of the 21 students were considered Extroverted, 14 Introverted, 11 Intuitive, 11 Sensing, 13 Feeling, 8 Thinking, 5 Perceiving, and 16 Judging.

In Fall 2023, 18 students were enrolled in Introduction to Biology Lab Section 1, and all were assigned to MBTI-based groups. Whereas, section 2, 16 students were enrolled, and all were assigned to LSB-based groups. For the MBTI population, in relevance to the ‘David Keirsey’s Temperament’, 5 students fell under the ‘Guardian’ category, 7 ‘Idealist’, 3 ‘Rational’, 3 ‘Artisan’. Based on their MBTI, 6 of the 18 students were considered Extroverted, 12 Introverted, 11 Intuitive, 7 Sensing, 13 Feeling. 5 Thinking, 10 Perceiving, and 8 Judging.

### Group Performance Metrics

Across all measured performance metrics, the MBTI based groups demonstrated consistently lower mean scores to the LSB organized groups **(Figure 3)**, indicating a broad decline across both interpersonal and affective dimensions of the learning environment. Mean scores for these metrics were uniformly lower in MBTI groups, with differences reaching statistical significance in each case (p<0.05), except for ‘contribution’. The largest effects were observed for belonging and satisfaction, where the MBTI-based cohort showed substantial decreases relative to the LSB-based cohort. Belonging exhibited a marked reduction (MBTI: 4.68 vs. LSB: 4.05, p = 0.0003), suggesting that MBTI-based grouping may have negatively impacted on students’ sense of inclusion and social integration. Satisfaction demonstrated the strongest effect overall (MBTI: 4.78 vs. LSB: 3.88, p < 0.0001), reflecting a pronounced decline in overall student experience within the MBTI-structured groups.

**Figure 2.**
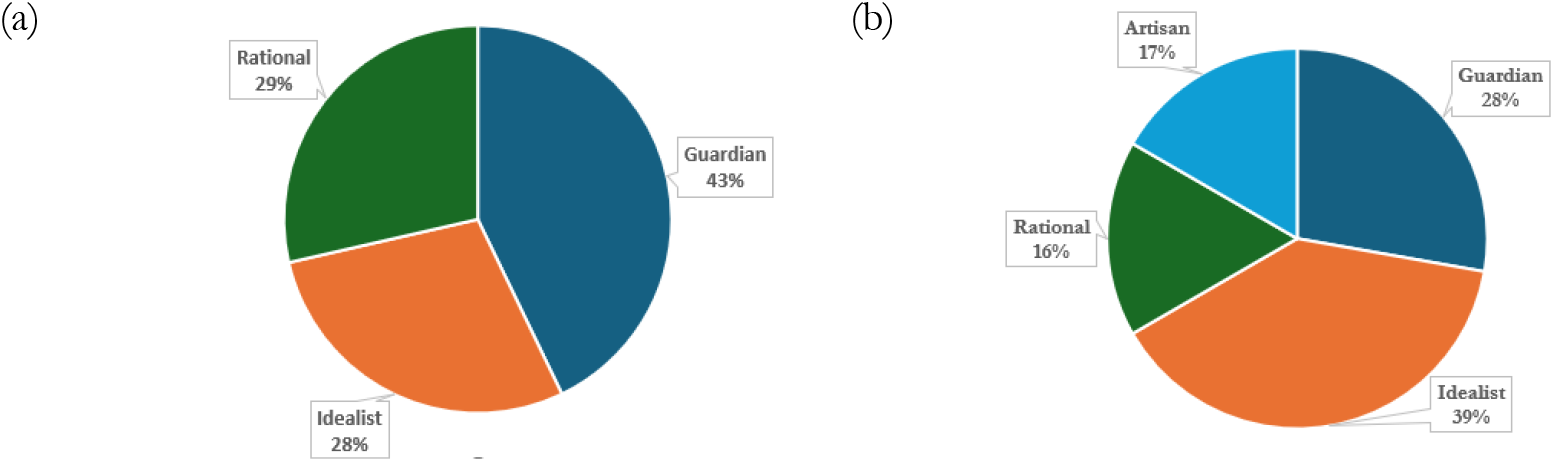
Keirsey’s Temperament Theory distribution of participants in MBTI-based groups. **(a)** Distribution of temperaments among students in the Cell Biology course (Fall 2023). **(b)** Distribution of temperaments among students in the Biology Laboratory Section 1. Keirsey’s theory classifies personalities into four temperaments reflecting behavioral patterns and decision-making styles: Artisans (SP) are adaptable and action-oriented; Guardians (SJ) are responsible and detail-focused; Idealists (NF) are empathetic and value-driven; and Rationals (NT) are strategic and logical. Each panel shows the proportion of participants in each temperament category.

**Figure 3.**
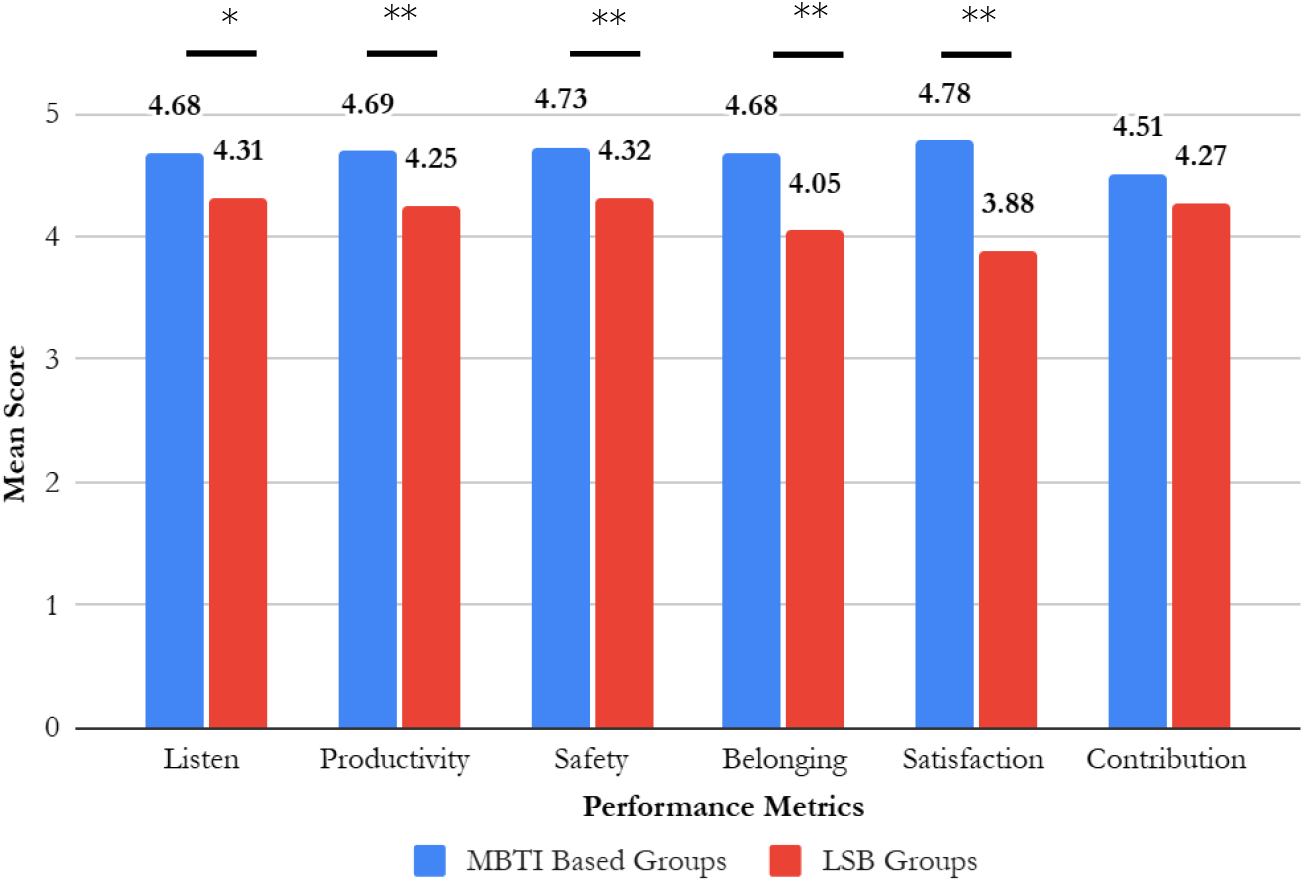
Comparison of Mean performance metrics between MBTI-based and LSB grouping strategies. The bar chart shows the mean scores for listening, productivity, sense of safety, belonging, satisfaction and contribution for students in Cell Biology 2023 (MBTI-based groups) and Cell Biology 2022 (LSB-groups). Asterisk indicates statistically significant differences between group strategies (*p < 0.05). All metrics showed significant decreases in the MBTI-based cohort except for contribution.

In contrast, contribution did not differ significantly between the two cohorts (MBTI: 4.51 vs. LSB: 4.27, p = 0.13), despite a numerically lower mean in the MBTI-based group.

### Role Fulfillment and Performance by Group Type

In the course wrapper, students were asked to rate role performance for themselves and their peers. Role performance ratings were calculated using responses from both self-assessments and peer assessments within each group. Mean scores were then computed for each role: Leader, Communicator, and Organizer, across both group types. Students in LSB groups scored significantly higher in the Leader and Communicator roles. In contrast, MBTI groups had higher scores for the Organizer role, although this difference was not statistically significant **(Figure 4)**.

**Figure 4.**
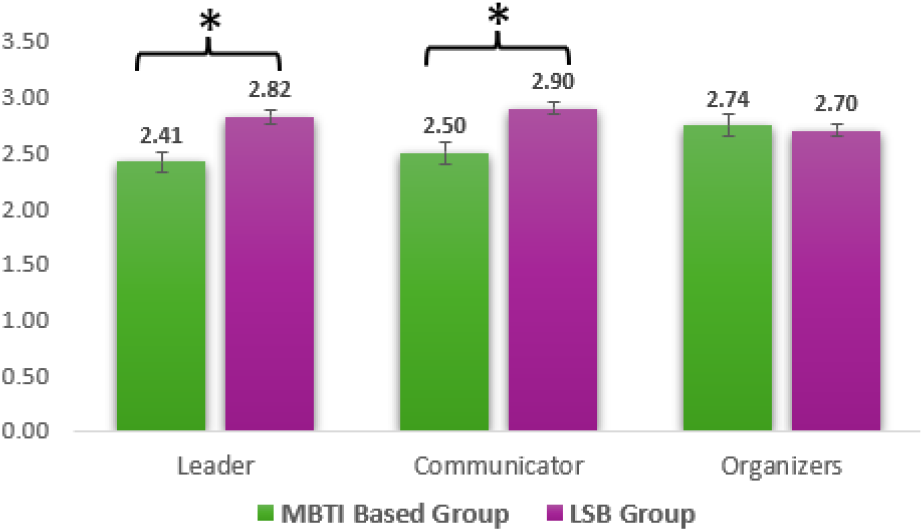
Role performance ratings by group type. Students in Biology Lab course, rated themselves and their group members on a 0–3 scale, where 0 indicated “never participated” and 3 indicated “exceeds expectations.” Mean scores were compared between MBTI-based and LSB groups using independent, pairwise t-tests for each role. LSB groups scored significantly higher in the Leader (t = 4.21, p < 0.001) and Communicator (t = 3.87, p = 0.002) roles. No significant difference was observed for the Organizer role (t = 0.45, p = 0.66). Asterisks on the figure indicate statistically significant differences between group types.

### Correlations Between Group Variables

In this study, data from two semesters of Cell Biology courses were used. Using data from *only* these two courses, Pearson’s correlation analysis was conducted to assess the relationship between group variables (e.g., satisfaction, productivity, and safety). Specifically, stronger correlations were observed in MBTI-Based Groups with respect to group satisfaction and productivity, while Learning Style-Based Groups showed notable correlations between safe-to-share and productivity **(Table 3)**. In MBTI-based groups, higher satisfaction ratings are correlated with higher perceived productivity, but with LSB groups higher satisfaction is correlated with the ‘safe to share’ variable.

**Table 3.** Correlation among group variables for LSB and MBTI-based groups. Shown are Pearson correlation coefficients between group satisfaction, productivity, and perceived safety to share opinions (Safe-to-Share) and the variables contribution, listening, productivity, safety, belonging, and satisfaction for two group types: Learning Style– Based (LSB) groups (N = 15; Fall 2022) and MBTI-Based groups (N = 21; Fall 2023) from a Cell Biology course. Correlation analyses were conducted using SPSS. Diagonal entries (---) indicate self-correlations. Asterisks denote statistical significance of the correlations (*p < 0.05; **p < 0.01)

| Variable | Contribution | Listen | Productive | Safe | Belonging | Satisfaction |
| --- | --- | --- | --- | --- | --- | --- |
| **LSB Group** |  |  |  |  |  |  |
| Satisfaction | 0.576* | 0.732** | 0.648** | 0.750** | 0.756** | --- |
| Productivity | 0.606* | 0.906** | --- | 0.933** | 0.843** | 0.648** |
| Safe to Share | 0.512 | 0.911** | 0.933** | --- | 0.954* | 0.750** |
| **MBTI Group** |  |  |  |  |  |  |
| Satisfaction | 0.387 | 0.644** | 0.568** | 0.523* | 0.739** | --- |
| Productivity | 0.362 | 0.593** | --- | 0.560** | 0.510* | 0.568** |
| Safe to Share | 0.511* | 0.678** | 0.560** | --- | 0.708** | 0.523* |

## DISCUSSION

The primary aim of this study was to compare MBTI-based and LSB grouping frameworks to determine which more effectively enhanced performance metrics and role fulfillment in Problem-Based Learning (PBL) environments. Across two course contexts: Cell Biology and Introduction to Biology Laboratory; LSB grouping consistently outperformed MBTI-based grouping on key performance metrics. In particular, **Figure 3** demonstrates that students in MBTI-based groups (Cell Biology 2023) reported lower scores across nearly all performance metrics of group functioning compared to students in LSB groups (Cell Biology 2022), including listening, productivity, sense of safety, belonging, and overall satisfaction. Contribution was the only metric that did not differ significantly between grouping strategies, suggesting a selective rather than uniform impact of grouping approach on student experiences.

LSB groups demonstrated significantly higher productivity scores in both Cell Biology and Biology Lab courses, reinforcing the idea that grouping students according to shared learning processes (e.g., active vs. reflective) may reduce friction in task execution and pacing. The additional findings related to listening and sense of safety further support this interpretation. Groups formed based on learning style alignment may facilitate more predictable communication patterns and mutual responsiveness, enabling students to feel heard and comfortable sharing ideas. In contrast, MBTI-based grouping, which intentionally promotes personality heterogeneity, may introduce variability in communication preferences and decision-making styles that complicate coordination, particularly in time-limited or task-intensive PBL settings.

Group satisfaction and belonging exhibited the largest and most robust differences between grouping strategies, with MBTI-based groups reporting markedly lower scores in both domains. These findings suggest that beyond task efficiency, learning-style homogeneity may play a critical role in fostering social cohesion and inclusion within small-group learning environments. Importantly, the absence of a significant difference in contribution indicates that students in MBTI-based groups did not perceive themselves as contributing less, despite experiencing lower satisfaction and weaker group climate.

Role performance analyses further clarify how grouping strategies shape group processes. As seen in **Figure 4**, Students in LSB groups scored significantly higher in Leader and Communicator roles, while no significant differences were observed for the Organizer role. Leadership and communication are inherently interactive and depend on shared expectations, pacing, and responsiveness. Learning-style similarity may facilitate smoother leadership emergence and clearer communication norms, whereas heterogeneous personality groupings may require greater negotiation and adaptation. The stability of the Organizer role across grouping conditions suggests that detail-oriented or task-structuring responsibilities may be less sensitive to group composition and more dependent on individual traits or clearly defined task demands.

Correlation analyses provide additional insight into the mechanisms underlying these differences. As seen in **Table 3** In MBTI-based groups, satisfaction was strongly correlated with perceived productivity, suggesting that these groups may be particularly sensitive to affective dynamics: when interpersonal functioning declines, productivity may be disproportionately affected. In contrast, LSB-based groups exhibited stronger associations between satisfaction and psychological safety, indicating that learning-style alignment may foster environments in which students feel comfortable expressing ideas, asking questions, and engaging in constructive disagreement. Sense of safety has been widely identified as a key predictor of effective teamwork and learning, particularly in collaborative educational contexts.

While MBTI-based grouping offers a structured framework for role differentiation and exposure to interpersonal diversity, the present findings suggest that learning style alignment may be a more robust predictor of effective group functioning in PBL settings. LSB grouping more directly supports productivity, satisfaction, and positive group climate. Importantly, these results do not imply that MBTI is pedagogically ineffective, but rather that its strengths may not align optimally with performance-driven PBL outcomes.

This study has several limitations. Sample sizes were modest and drawn from a single institution, limiting generalizability. The Cell Biology grouping conditions were implemented across different semesters rather than within the same cohort, introducing potential cohort effects. Additionally, outcome measures relied on self-reported Likert-scale instruments, which may be subject to response bias. Future studies should incorporate larger, multi-institutional samples, randomized within-cohort designs, and objective performance measures to further clarify how grouping strategies influence learning outcomes.

## CONCLUSIONS

This study demonstrated that LSB grouping, grounded in Kolb’s Experiential Learning Theory, is more effective than MBTI-based grouping in promoting group productivity, satisfaction, and role fulfillment in undergraduate PBL settings. While MBTI-based groups showed meaningful internal correlations between satisfaction and productivity, they consistently underperformed relative to LSB groups on primary outcome measures. These findings suggest that how students learn may be more consequential for group effectiveness than how they interact personality-wise. For instructors designing collaborative learning environments, particularly in science courses that rely heavily on teamwork, grouping students by learning style may offer a practical and evidence-based strategy for enhancing both performance and student experience.

## ACKNOWLEDGEMENTS

We thank all the students who participated in the surveys for their time and contributions to this study. We also acknowledge the Undergraduate Research Student-Centered Activities (URSCA) program at the University of Wisconsin–Superior for funding this research. Additionally, we appreciate the support of course instructors and teaching staff who facilitated participant access and assisted with logistical aspects of data collection.

## ABOUT THE STUDENT AUTHOR

Basith Nasik conducted this research as an undergraduate student at the University of Wisconsin-Superior graduating with a Bachelor of Science in Biology (Pre-Health concentration) & Biochemistry in May 2024. Currently as a graduate student in the Biology Department at the University of Wisconsin-Madison, he focuses on the Hippo cell proliferation pathway and protein phosphorylation. He aspires to contribute to translational research, integrating interdisciplinary approaches to address complex scientific questions, and aims to advance both knowledge and practical applications that can positively impact society.

## PRESS SUMMARY

Group work is a cornerstone of many college science courses, yet how students are assigned to teams is often overlooked. In this study, we found that forming groups based on students’ learning styles led to higher productivity, stronger leadership and communication, and greater overall satisfaction compared to grouping students by personality type using the Myers-Briggs Type Indicator (MBTI). Students in learning style–based groups reported feeling more comfortable sharing ideas and worked more efficiently together, while personality-based groups showed lower satisfaction and performance overall. These findings suggest that aligning how students learn may be more important than mixing personality types when designing effective collaborative learning experiences in higher education.

## REFERENCES

1. Knowledge, Developing Transferable. “EDUCATION FOR LIFE and WORK.” (2012) Pérez Zúñiga, Ricardo, M. A. R. I. O. MARTíNEZ GARCíA, and Francisco Eduardo Oliva Ibarra. “Employability and its relationship with the competency-based approach, teaching methodologies, and assessment in higher education: a systematic review.” Frontiers in Education. Vol. 10. Frontiers.

2. Pandya, Bharti, Umar Ruhi, and Louise Patterson. “Preparing the future workforce for 2030: the role of higher education institutions.” Frontiers in Education. Vol. 8. Frontiers Media SA, 2023.

3. Romanova, Anna. “Personal Belbin types and MBTI preferences combination in group work assuring success.” (2018).

4. Oflaz, Merve, and Turgut Turunc. “The effect of learning styles on group work activities.” Procedia-Social and Behavioral Sciences 46 (2012): 1333–1338.

5. Myers, Briggs. “Katherine Cook Briggs and Isabel.” Key Thinkers in Individual Differences: Ideas on Personality and Intelligence (2019).

6. Kolb, David A. Experiential learning: Experience as the source of learning and development. FT press, 2014.

7. Fleming, Neil D., and Colleen Mills. “Not another inventory, rather a catalyst for reflection.” To improve the academy 11.1 (1992): 137–155.

8. Xu, L. Development of a visual complex for MBTI: qualification thesis 022 Design. Kyiv: KNUTD (2024). 70 p.

9. Ullah, A., Uddin, F., Khan, S., Imran, M. (2024). Exploring the impact of MBTI personality types on teaching methods. Qlantic J. Soc. Sci. 5(3), 309–323.

10. Sudria, I.B.N., et al. (2018). Effect of Kolb’s learning styles under inductive guided-inquiry learning on learning outcomes. Int. J. Instr. 11(1), 89–102.

11. Lawrence, W.K. (2015). Learning and personality: The experience of introverted reflective learners in a world of extroverts. Cambridge: Cambridge Scholars Publishing.

12. Louis, G. Personality type and leadership dynamics: Exploring MBTI’s influence on student leader’s academic performance, work-life balance, and stress management.

13. Barrows, H.S., Tamblyn, R.M. (1980). Problem-based learning: An approach to medical education. New York: Springer Publishing Company.

14. Moreland, R.L., Levine, J.M., Wingert, M.L. (2018). Creating the ideal group: Composition effects at work. In: Understanding group behavior. London: Psychology Press, 11–35.

15. Sanders, D. (2014). Personality type, leadership, collaborative decision making and collective efficacy. Diss. Oakland University.

16. De Raad, Boele. The big five personality factors: the psycholexical approach to personality. Hogrefe & Huber Publishers, 2000.

17. Stein, R., Swan, A.B. (2019). Evaluating the validity of Myers-Briggs Type Indicator theory: A teaching tool and window into intuitive psychology. Soc. Pers. Psychol. Compass 13(2), e12434.

18. Ph’ng, L.M., Thang, S.M., Nambiar, R.M.K. (2015). Matching teaching styles and learning styles: What happens in the case of a mismatch? e-BANGI 10(1), 66.

19. Keirsey, David, and Marilyn M. Bates. Please understand me: Character & temperament types. Del Mar, CA: Prometheus Nemesis Book Company, 1984.

20. Sanders, Dyanne. Personality type, leadership, collaborative decision making and collective efficacy. Diss. Oakland University, 2014.

